# Protein crosslinking as a therapeutic strategy for SOD1-related ALS

**DOI:** 10.1101/2021.06.23.449516

**Authors:** Md Amin Hossain, Richa Sarin, Daniel P. Donnelly, Brandon C. Miller, Joseph P. Salisbury, Jeremy B. Conway, Samantha Watson, Jenifer N. Winters, Novera Alam, Durgalakshmi Sivasankar, Aparna C. Ponmudiyan, Tanvi Gawde, Sunanda Kannapadi, Jared R. Auclair, Lee Makowski, Gregory A. Petsko, Dagmar Ringe, David J. Greenblatt, Mary Jo Ondrechen, Yunqiu Chen, Roman Manetsch, Jeffrey N. Agar

**Affiliations:** Department of Chemistry and Chemical Biology, Northeastern University, Boston, MA 02115; Barnett Institute of Chemical and Biological Analysis, Boston, MA 02115; Biogen Inc, Cambridge, MA 02142; Department of Bioengineering, Northeastern University, Boston, MA 02115; Ann Romney Center for Neurologic Diseases at Brigham and Women’s Hospital, Harvard Medical School, Boston, MA 02115; Departments of Chemistry and Biochemistry, and Rosenstiel Center for Basic Medical Research, Brandeis University, Waltham, MA 02454; School of Medicine, Tufts University, Boston, MA 02111; Department of Pharmaceutical Sciences, Northeastern University, Boston, MA 02115.

**Keywords:** SOD1, Amyotrophic Lateral Sclerosis (ALS), pharmacological chaperone, cross-linking, stabilization

## Abstract

Mutations in the gene encoding Cu-Zn superoxide dismutase 1 (SOD1) cause a subset of familial amyotrophic lateral sclerosis (fALS). One effect of these mutations is that SOD1, which is normally a stable dimer, dissociates into toxic monomers. Considerable research efforts have been devoted to developing compounds that stabilize the dimer of fALS SOD1 variants, but these have not yet resulted in an approved drug. We demonstrate that a cyclic thiosulfinate cross-linker can stabilize prevalent disease-causing SOD1 variants. The degree of stabilization afforded by cyclic thiosulfinates (up to 24 °C) is unprecedented. We show this compound works rapidly in vivo with a half-life of ~3 days. The efficacy, low toxicity, and pharmacodynamics of cross-linker mediated stabilization make it a promising therapeutic approach for SOD1-related fALS.

**Significance statement:** Cyclic thiosulfinate *S*-XL6 enables the kinetic stabilization of ALS-associated SOD1 variants, in vivo.

## Introduction

ALS is a fatal neurodegenerative disease that often leads to death within 2-5 years of diagnosis (1, 2). 5%-10% of ALS cases are familial while the rest are sporadic (3, 4). Over 180 SOD1 mutations in the gene encoding SOD1 contribute to approximately 20% of fALS cases (5, 6). These mutations are dominantly inherited, highly penetrant, and associated with varying degrees of disease severity and loss of enzymatic activity (7, 8). For example, patients harboring the SOD1 variant, SOD1^A4V^ tend to survive one year after diagnosis, whereas those with SOD1^H46R^ average 18 years after diagnosis (9). A number of studies have implicated *wild-type* SOD1 in sporadic ALS. For example, post-translational modification (PTMs) such as oxidation of Cys111 (10) are toxic and autoantibodies (11) to SOD1 are more prevalent. The role of *wild-type* SOD1 in ALS remains a subject of debate (12).

*Wild-type* SOD1 is a dimer. A common property of ALS-associated SOD1 mutations is that they increase the propensity for the dissociation of the SOD1 dimer (13). Small molecules are therefore being developed to stabilize the quaternary structure of fALS SOD1 variants (14, 15). An analogous strategy was employed successfully for tafamidis, a drug for transthyretin amyloidosis and transthyretin cardiomyopathy, which stabilizes the native transthyretin tetramer (16). However, this strategy has not yet resulted in a treatment for SOD1-related fALS. For example, molecules intended to stabilize SOD1 via non-covalent interactions at the dimer interface (17) instead bound to the β-barrel region of the SOD1 aggregates (14) and often exhibited plasma protein binding (15).

We are pursuing an alternative approach to stabilize the SOD1 dimer. We postulated that two free cysteines (a sulfhydryl, i.e., not involved in a disulfide) situated on adjacent monomers, Cys111_a_ and Cys111_b_ (within 9-13 Å of each other, depending upon Cys residue orientation), could be cross-linked to prevent dimer dissociation. Cysteine residues can be targeted with high selectivity due to the unique reactivity (nucleophilicity and polarizability) of the thiolate functional group. This is evidenced by numerous drugs that form covalent bonds to cysteine residues, including the irreversible aldehyde dehydrogenase (ALDH1A1) inhibitor disulfiram (Antabuse) (18); the blockbuster proton pump inhibitors, e.g., omeprazole (Prilosec) and its single enantiomer esomeprazole (Nexium) (19); and second-generation kinase inhibitors, e.g., afatinib (Gilotrif) (20). In proof-of-concept experiments, we demonstrated that tethering the Cys111 pair on the disease variants SOD1^G93A^ and SOD1^G85R^ could stabilize the SOD1 dimer and rescue superoxide dismutase activity (21). The bifunctional maleimides used in these studies, however, are toxic (22) because they target any exposed free cysteine, many of which serve essential catalytic and redox roles for other proteins. The molecule ebselen, which stabilizes fALS SOD1 variants by binding Cys111 (23) could exhibit a similar liability, as evidenced by its off-target binding to Cys6 (24). Therefore, a therapeutically viable cross-linking mechanism requires the ability to bind the Cys111 pair while avoiding other cysteines.

We recently introduced cyclic thiosulfinates, which selectively cross-link pairs of free cysteines, while avoiding “dead-end” modifications (i.e., not resulting in a cross-link) of lone cysteines (25). Cyclic thiosulfinates are one oxygen (*S*-oxo) derivatives of cyclic disulfides. Well-known and well-tolerated cyclic disulfides include the natural product asparagusic acid and the multi-enzyme cofactor and dietary supplement α-lipoic acid. The cyclic thiosulfinate 1,2-dithiane-1-oxide (sulfur cross-linking 6-membered ring, hereafter *S*-XL6, **Fig. 1**) and β-lipoic acid (the *S*-oxo derivative of α-lipoic acid), (25) target the Cys111_a_ and Cys111_b_ residues on adjacent SOD1 monomers, forming a disulfide bond with each cysteine thiolate. To test the efficacy of *S*-XL6, we have chosen disease variants ranging from metallated, fully active (SOD1^A4V^ and SOD1^G93A^) to mostly inactive with impaired metal binding (SOD1^H46R^ and SOD1^G85R^). These same variants also span a wide range of prognoses (1 year to 18 years survival). In this study, we demonstrate *S*-XL6-mediated dimer stabilization of these fALS SOD1 variants; assess the impact of the cross-linker on the thermal stability and structure; test in cellulo activity and toxicity, and demonstrate target engagement in an ALS mouse model.

**Fig. 1.**
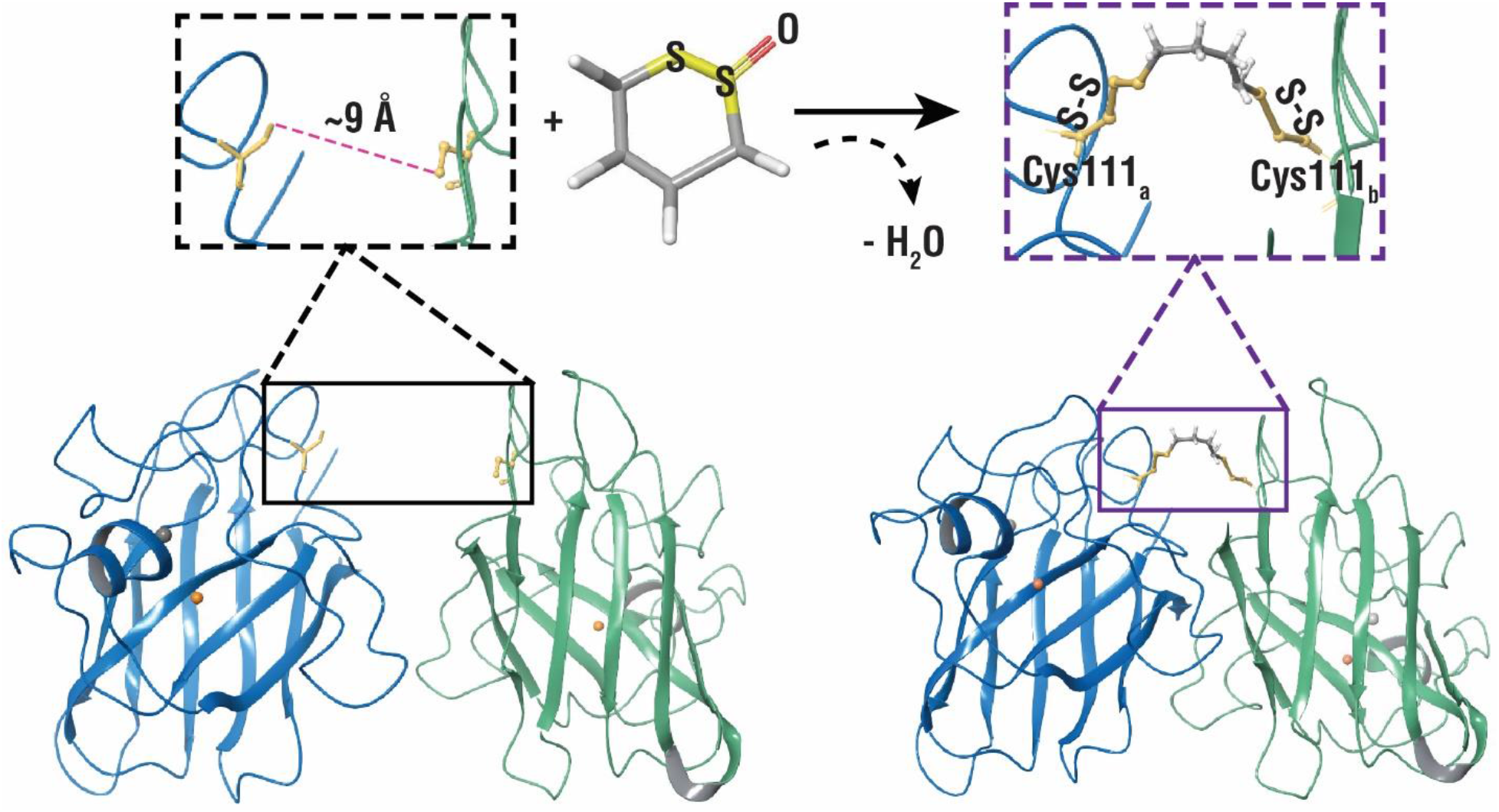
Cyclic thiosulfinate *S*-XL6 cross-links SOD1 variants via Cys111 residues on adjacent monomers. Crystal structure of *wild-type* SOD1 (PDB ID: 1SPD, cartoon representation generated with Maestro 11.8) highlighting opposing Cys111 residues on both monomer A (blue) and monomer B (green) with a representation of the *S*-XL6 cross-linked SOD1 dimer. The cross-linking reaction proceeds through an initial thiolate-disulfide interchange between the cyclic thiosulfinate and a cysteine thiolate, generating a sulfenic acid intermediate due to the opening of the ring structure. This sulfenic acid intermediate forms a cross-link by rapid condensation with a second cysteine thiolate. Cu and Zn molecules are represented by orange and grey spheres, respectively.

## Results

### S-XL6 cross-links and stabilizes fALS SOD1 variants

We employed a mass spectrometry (MS) assay for intact protein crosslinking (21). In this assay, any non-covalent (i.e., native) dimer is dissociated by a combination of acidic, organic media and MS source conditions. Using this assay, we observed *S*-XL6-dependent formation of cross-linked dimers for *wild-type* SOD1, SOD1^A4V^, SOD1^G93A^, SOD1^H46R^, and SOD1^G85R^ (**Fig. 2)**. This reaction was efficient (full conversion of monomer to dimer) and stoichiometric (occurring at ratios as low as 1:1 cross-linker to dimer). Dead-end modifications of SOD1 were not observed in these assays. We confirmed the mechanism of action (MoA, e.g., loss of oxygen from the cross-linker concomitant with dimer formation, **Fig. 1**) and determined the site of cross-linking using endoproteinase digestion and MALDI (Matrix-Assisted Laser Desorption Ionization) peptide mass fingerprinting analysis of cross-linked samples (**Fig. S1**).

**Fig. 2.**
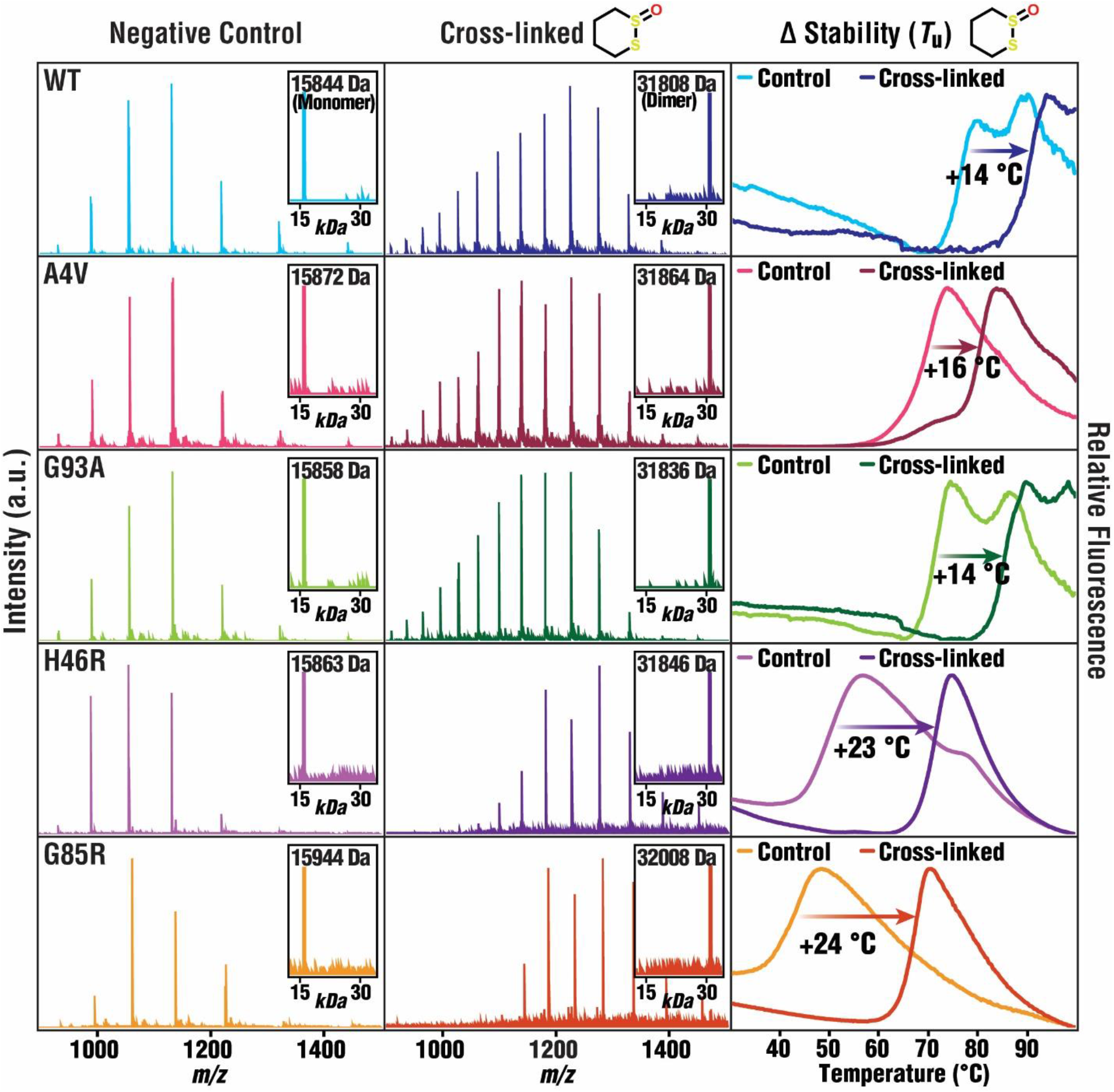
*S*-XL6-mediated cross-linking and stabilization of *wild-type* and fALS SOD1 variants. Mass spectra of untreated (**left column,** average mass [± 1 Da]) and *S*-XL6 cross-linked (**middle column,** average mass [± 2 Da]) proteins are consistent with cross-linking. DSF results (**right column**) indicate that cross-linking increased the thermal stability of SOD1 and its variants. Note: two inflections can be observed when there is a mixture of partially (lower Δ*T*_u_) and fully metallated (higher Δ*T*_u_) SOD1 proteoforms, which applies to the following proteins: untreated *wild-type* SOD1 unfolds at 75.9 °C and 87.6 °C; untreated SOD1^G93A^ unfolds at 70.6 °C and 84.4 °C. We quantify Δ*T*_u_ as the difference in unfolding temperature measured between the major inflections of the untreated and cross-linked samples. The unfolding temperature of the untreated and *S*-XL6 cross-linked SOD1 proteins are as follows: *wild-type* (WT) from 75.9 °C to 90.2 °C (Δ*T*_u_ ~14 °C); SOD1^A4V^ from 62.9 °C to 79.3 °C (Δ*T*_u_ ~16 °C); SOD1^G93A^ from 70.6 °C to 85.0 °C (Δ*T*_u_ ~14 °C); SOD1^H46R^ from 46.3 °C to 69.3 °C (Δ*T*_u_ ~23 °C); SOD1^G85R^ from 40.8 °C to 65.1 °C (Δ*T*_u_ ~24 °C). All samples were analyzed in triplicate (standard errors < 0.3 °C).

Mutations in SOD1 that cause ALS destabilize the native dimer to varying degrees. Loss of stability of fALS variants correlates with disease severity, specifically with rapid progression (26). To understand how cross-linking effects SOD1 stability, we quantified changes in unfolding temperature (Δ*T*_u_, hereafter thermal stability). Purified *wild-type*, SOD1^A4V^, SOD1^G93A^, SOD1^H46R^, and SOD1^G85R^ proteins were treated with *S*-XL6. Thermally-induced unfolding of SOD1 variants was measured using a previously published method, Differential Scanning Fluorimetry (DSF) (**Fig. 2, right**) (27). The increases in thermal stability attained with our compounds (14-24 °C) compare favorably with that of other preclinical candidates for stabilizing SOD1 (e.g., 11 °C ebselen analogues) (28), and with the only approved drug with a kinetic stabilizing mechanism, tafamidis (6 °C) (29).

### S-XL6 promotes SOD1 dimer formation *in cellulo*

We characterized the effectiveness of our crosslinker using Hep G2 cells. SOD1 cross-linking was monitored using SDS-PAGE and western blotting. *S*-XL6 cross-linked *wild-type* SOD1 in cells in both PBS buffer and serum. The half maximal effective concentrations (EC_50_) were circa (*ca*.) 5 μM and 10 μM (**Fig. 3*A***), respectively, consistent with minor association with serum proteins. The minimal plasma protein binding is promising, and is enabled by cyclic thiosulfinates’ ability to avoid lone cysteines, including the highly abundant lone cysteine (Cys34) (30) of serum albumin. These results are also consistent with previous research indicating that cyclic thiosulfinates are actively transported across the cellular membrane (25) where they remain intact (31).

**Fig. 3.**
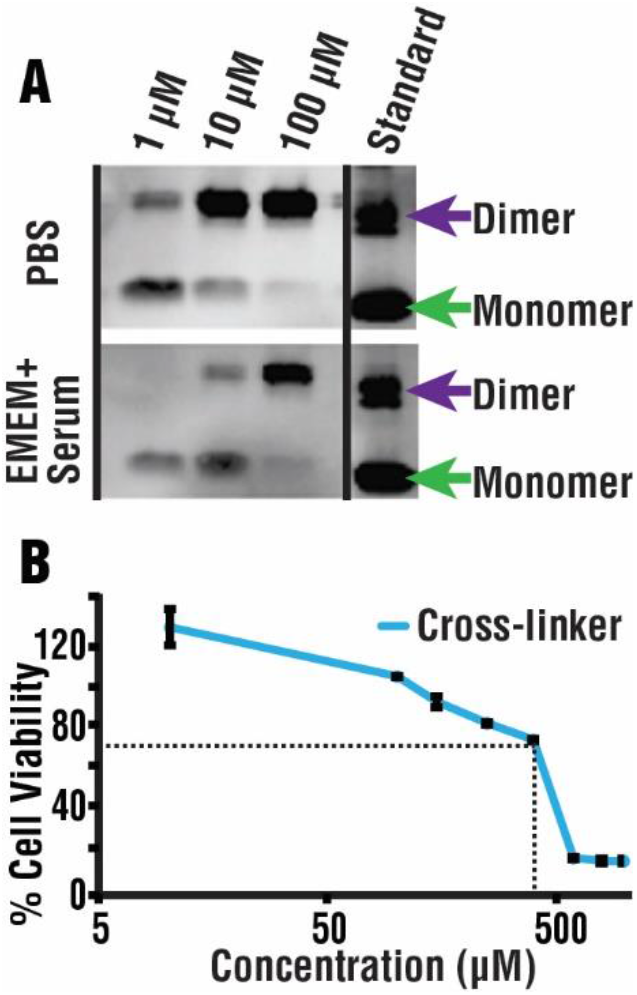
Cross-linking at sub-toxic concentrations within cells. (****A****) Western blot analysis using a SOD1-selective antibody indicates that cross-linking proceeds in HEP G2 cells with an EC_50_ of ~5 μM (in PBS buffer) or 10 μM (in complete growth media, including 10% serum). (****B****) Low toxicity of *S*-XL6 (LC_50_ ~446 μM) was observed using the MTT (3-(4,5-Dimethylthiazol 2-yl)-2,5-diphenyltetrazolium bromide) cytotoxicity assay (triplicate analysis). Results are shown as percentage of viable cells compared to a vehicle control. Note: cell viability <70% (dotted line) is generally considered as the threshold for cytotoxicity, and viabilities >100% indicate a trophic effect. Controls included EMEM with 0.1% DMSO (dimethyl sulfoxide, negative control) and 500 μM of chlorpromazine (positive control).

The cytotoxicity of *S*-XL6 was measured in Hep G2 using a standard MTT assay (LC_50_ ~446 μM, **Fig. 3***B*). Given that the LC_50_ is 90x the EC_50_, *S*-XL6 can promote dimer formation *in cellulo* with minimal toxicity. This warranted further exploration of *S*-XL6 as a potential preclinical candidate, *vide infra*.

### Target stabilization and pharmacodynamic profiling

To assess the time course of *S*-XL6 target engagement in vivo, pharmacodynamic (PD) profiling was carried out using mice expressing human fALS variant, namely the well-established “fast-line” B6SJL-Tg(SOD1*G93A)1Gur/J (13). Mice were dosed with *S*-XL6 via intravenous (IV) injection and blood was collected at different time points, post dose, from the tail vein. To detect the intact SOD1^G93A^ protein-cross-linker complex, we utilized a facile MS assay, namely a combination of solvent-extraction, hemoglobin precipitation, and liquid chromatography coupled with mass spectrometry (LC-MS) (32). We observed that a single IV dose of *S*-XL6 at 10 mg/kg converted 63% of the SOD1^G93A^ (**Fig. 4*D***) into a cross-linked dimer at 1-hour post-dose. A notable half-life of *ca.* 68 hrs. was observed (**Fig. 4*B***). The accuracy of the MS assay (**Fig. 4**) allows us to confirm the following aspects of our proposed MoA (**Fig. 1**), in vivo: cyclic thiosulfinate’s oxygen is lost upon cross-linking, and only one cross-linker binds per SOD1 dimer.

**Fig. 4.**
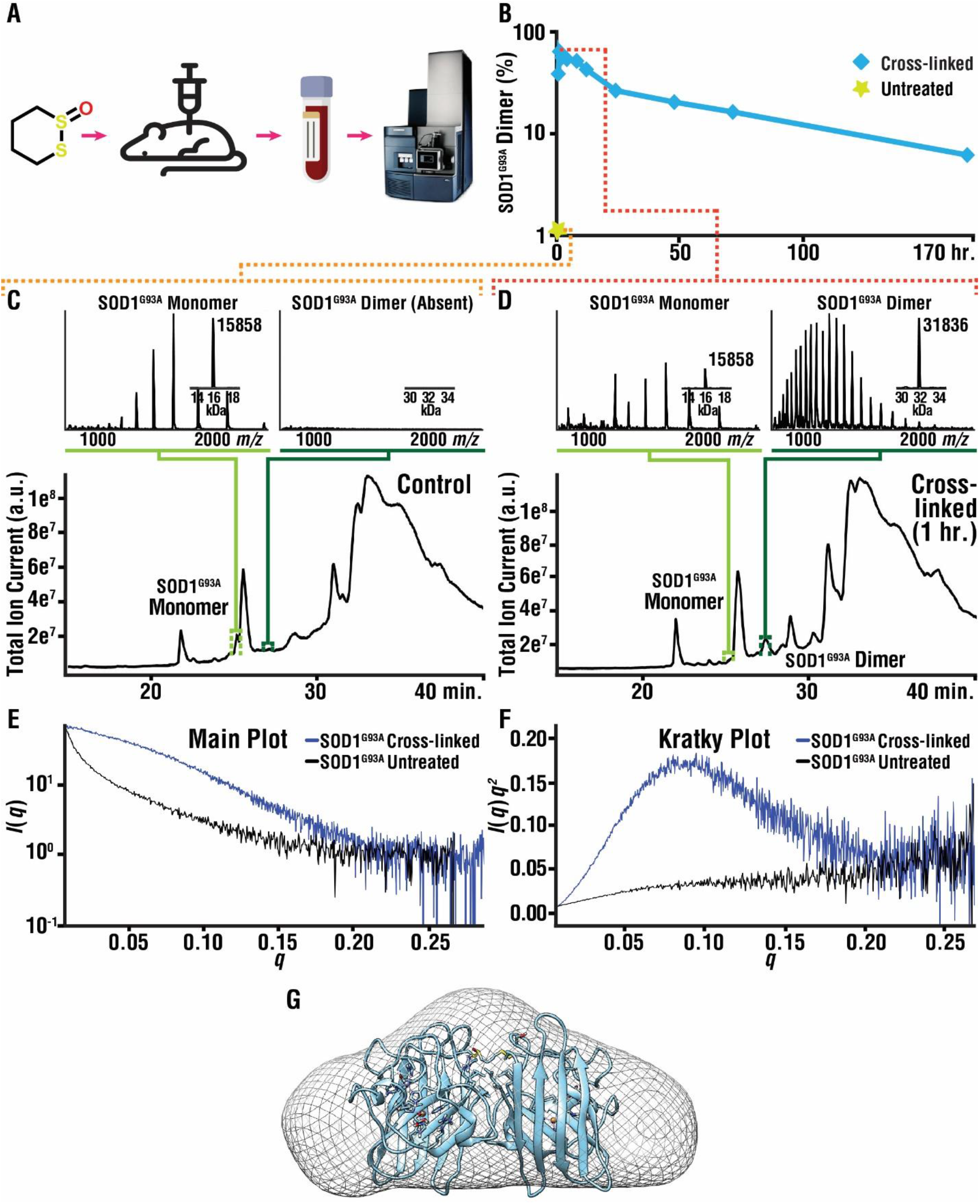
*S*-XL6 cross-links SOD1 in an ALS mouse model & results in a more folded SOD1 dimer. (***A***) Schematic of the pharmacodynamic workflow. (*B*) Pharmacodynamic profiling of red blood cell (RBC) proteins by LC-MS analysis indicates the formation of an *S*-XL6 cross-linked dimer in treated SOD1^G93A^ mice. Hemizygous fALS SOD1^G93A^ mice were dosed once at 10 mg/kg with *S*-XL6 via tail vein injection, blood was collected periodically over a 7-day period, and LC-MS analysis (***C* and *D***) was performed to assess percentage of cross-linked SOD1 (details in **Table S1**). Spectra represent a 6 second average at peak apex. The half-life of cross-linked SOD1^G93A^ dimer was determined to be *ca*. 68-hours. (***E* and *F***) In vitro SAXS experiments for SOD1^G93A^: Semi-log plot of the scattering intensity (I, log scale) as a function of momentum transfer, *q*; and Kratky plot [*q^2^*•*I*(*q*) versus *q*]. Plot shape for untreated SOD1^G93A^ is consistent with an unfolded protein, whereas plot shape for cross-linked SOD1^G93A^ is consistent with a folded, globular structure. The estimated radius of gyration is ~20 Å. (****G****) A 3-D reconstruction of cross-linked SOD1^G93A^ is superimposed upon the SOD1^G93A^ crystal structure (PDB: 3GZO).

Previous studies have shown that Small Angle X-Ray Scattering (SAXS) can monitor changes to protein tertiary and quaternary structure in solution. Therefore, SAXS (**Fig. 4 *E* and** *F*) was performed to determine the effects of cross-linking upon the structure of fALS variant SOD1^G93A^, the same variant expressed by the mice that were dosed in the present study. For SAXS studies, SOD1^G93A^ was purified from yeast because the milligram protein quantities required could not be obtained from mice. Whereas previous SAXS studies of SOD1^G93A^ preparations showed a radius of gyration (*R*g) of 20 to 24 Å (33, 34), our SAXS data for as-isolated SOD1^G93A^ were consistent with a relatively unfolded (possibly aggregating) SOD1 structure and could not be fit to determine an *R*g. The *R*g of 20 Å estimated from scattering data was consistent with *S*-XL6 treated SOD1^G93A^ being a native dimer.

### Higher order structural analysis of cross-linked SOD1

To determine the impact of cross-linking on the structure of fALS SOD1 variants, we utilized hydrogen deuterium exchange mass spectrometry (H/D-X MS). H/D-X MS assesses structure and dynamics by measuring differences in deuterium uptake (35). Following cross-linking, we observed that the structures of fALS SOD1 variants bear a closer resemblance, but do not fully recapitulate, the *wild-type* SOD1 structure (**Fig. 5**). For example, decreased uptake at the N- and C-termini of SOD1 (the dimer interface) was observed in all cross-linked variants. In all but one variant, SOD1^G93A^, the differences in uptake around residues 37-43 more closely resembled *wild-type*. Notably, structural changes in this region are responsible for the “gain of interaction” with the disordered electrostatic loop, which has been proposed to lead to the aggregation of fALS variants (36).

**Fig. 5.**
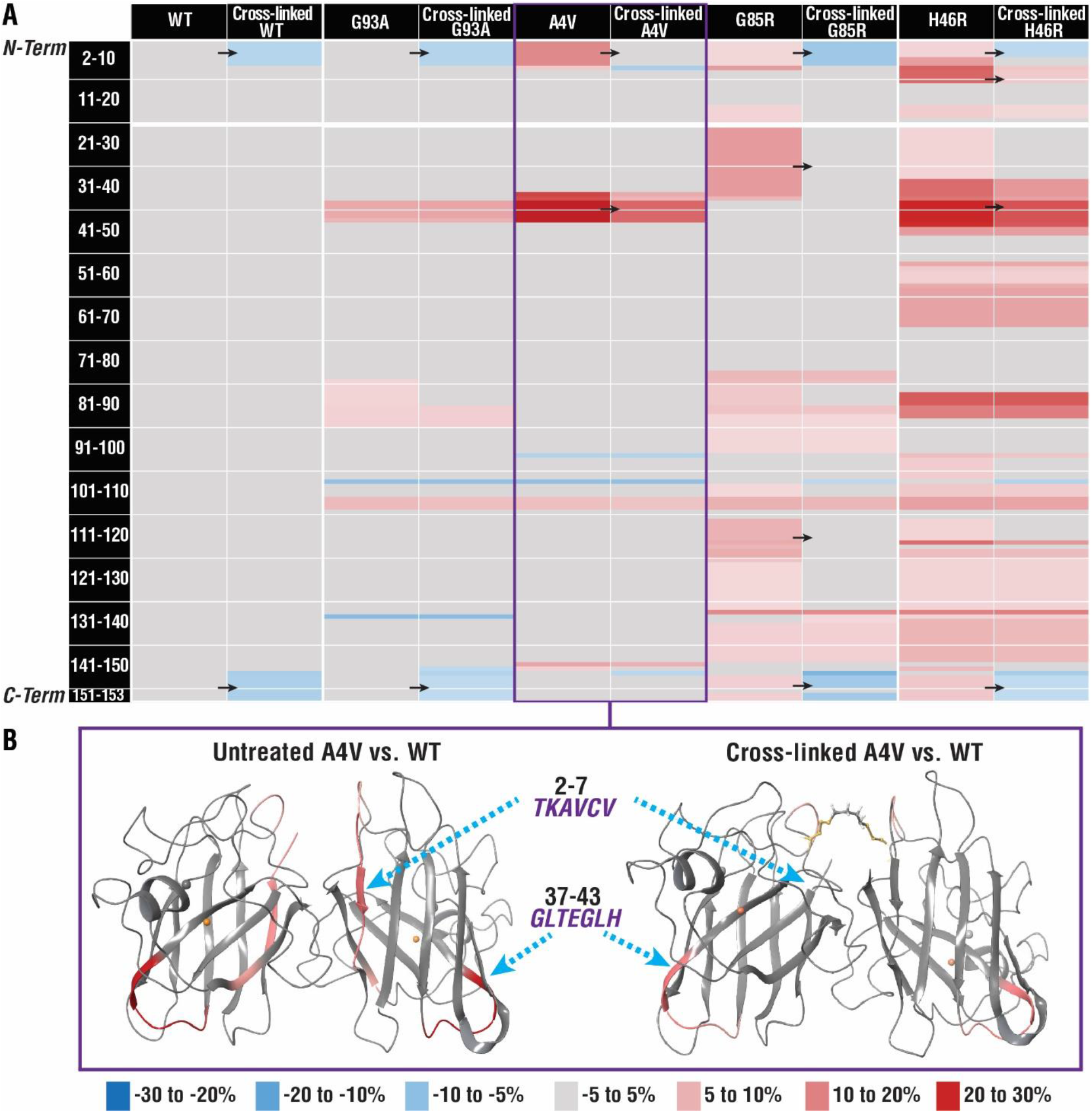
Cross-linking enhances the structure of fALS SOD1 variants. Differences in deuterium uptake (ΔU, legend shown below part b) of untreated and cross-linked variants for the 4-hour timepoint compared to the *wild-type* SOD1 (WT) protein are reported here. (****A****) → indicates areas where prominent shifts in ΔU were observed (e.g., residues 2-7 for SOD1^A4V^ untreated 13.8% to 1.2% cross-linked, for full results see **Fig. S2**). (****B****) ΔU for untreated and cross-linked SOD1^A4V^ mapped onto the cartoon representation of *wild-type* SOD1 structure (PDB ID: 1SPD), generated with Maestro 11.8. ΔU for all timepoints (15s, 50s, 500s, 1 hour, 4 hours) are reported in **Fig. S2**.

Previous studies have shown that fALS variants can be broadly classified as enzymatically active “*wild-type*-like variants” (e.g., SOD1^A4V^ and SOD1^G93A^), and as inactive, less-folded, predominantly monomeric (36, 37), “metal-deficient” variants (e.g., SOD1^G85R^ and SOD1^H46R^) (38, 39). Consistent with this, larger differences in uptake via H/D-X MS were observed for SOD1^G85R^ and SOD1^H46R^. Subtle (<5%) differences in deuteration levels were also observed in other regions of the SOD1 variants. In summary, cross-linking makes the structure and dynamics of fALS SOD1 variants more similar to that of *wild-type* SOD1, and in a manner that is consistent with reducing their aggregation propensity.

## Discussion

We have shown that a cyclic thiosulfinate cross-linker, *S*-XL6, cross-links and stabilizes SOD1 fALS variants with diverse physiochemical properties and disease severities. These variants included the prevalent G85R and H46R SOD1 variants, which provided a major challenge for our dimer cross-linking approach because they exist in a predominantly monomeric state. Notably, the ~68 hour biological half-life of the *S*-XL6 cross-linked complex is almost seven times higher than previously reported studies (e.g., 10 hour half-life of SOD1^G93A^) (40). DSF and SAXS results indicated that cross-linking increased stability and H/D-X results indicated that cross-linking should decrease the aggregation propensity of fALS variants. These results are promising given that structural instability and higher aggregation propensity are risk factors for SOD1-related fALS(9, 41), but are only the first step in therapy development. Additional preclinical studies e.g., addressing selectivity and Drug Metabolism and Pharmacokinetics (DMPK) are underway. The focus of this study was the kinetic stabilization of SOD1 dimer. The most effective preclinical therapeutic strategy for SOD1 related-fALS has been the stabilization of SOD1 via the CuATSM (Cu (II)-diacetyl-bis(N(4)-methylthiosemicarbazone)-mediated incorporation of metals (42). We anticipate there would be a benefit to combining these approaches.

## Materials and Methods

### Synthesis of cyclic thiosulfinate 1,2-dithiane-1-oxide (*S*-XL6)

Synthesis of 1,2-dithiane-1-oxide (*S*-XL6) was achieved according to the previously published literature procedure (25). To a round-bottom flask was added I_2_ (2.08 g, 8.18 mmol, 0.20 equiv.) and DMSO (2.91 mL, 40.90 mmol, 1.0 equiv.). The resulting dark brown solution was stirred gently. 1,4-butanedithiol (4.76 mL, 40.90 mmol, 1.0 equiv.) was dissolved in CH_2_Cl_2_ (16.3 mL) and added dropwise to the stirring solution of I_2_ in DMSO. The rate of stirring was increased gradually with the addition of 1,4-butanedithiol in CH_2_Cl_2_. Upon completion of addition the resulting solution was stirred for 1 h at 23 °C. The light brown solution was quenched with slow addition of 10% aq. Na_2_S_2_O_3_ (25 mL). The layers were separated, and the aqueous layer was extracted with CH_2_Cl_2_ (3 × 50 mL). The combined organic layers were concentrated under reduced pressure. The concentrated light-yellow oil was taken up in 100 mL EtOAc and washed with sat. aq. NaCl (50 mL). The layers were separated, and the aqueous layer was extracted with EtOAc (3 × 75 mL). The combined organic layers were dried over Na_2_SO_4_ and concentrated under reduced pressure to give pure 1,2-dithiane (4.82 g, 98% yield) as a faint yellow solid. Rf = 0.80 (5:1 hexanes:EtOAc). 1,2-dithiane was then used without further purification to form 1,2-dithiane-1-oxide (*S*-XL6).

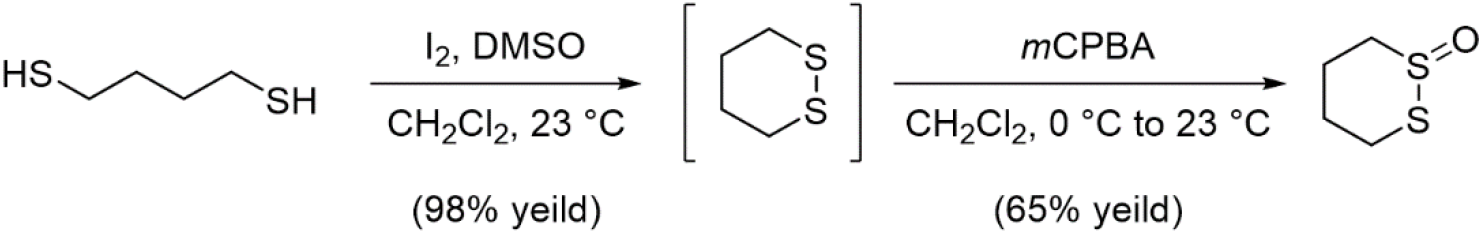

In a round-bottom flask, 1,2-dithiane (4.82 g, 40.09 mmol, 1.0 equiv.) was dissolved in CH_2_Cl_2_ (40.0 mL) and cooled to 0 °C. A solution of *m*CPBA (9.48 g, 73% weight, 40.09 mmol, 1.0 equiv) in CH_2_Cl_2_ (100.0 mL) was added dropwise via addition funnel. The resulting colorless, cloudy solution was allowed to stir at 0 °C for 1 hour while warming to 23 °C and allowed to stir 1 hour at 23 °C. The reaction was cooled again to 0 °C and quenched with solid Na_2_CO_3_ (40.0 g, 377 mmol, 1 g per mmol *m*CPBA). The resulting dense slurry was filtered through a short plug of celite and concentrated under reduced pressure. The resulting cloudy oil was purified by flash column chromatography on silica gel with 2% MeOH / CH_2_Cl_2_ to give *S*-XL6 (3.53 g, 64% yield) as a colorless solid. Rf = 0.33 (2% MeOH / CH_2_Cl_2_). ^1^H NMR (500 MHz, CDCl_3_, δ): 1.82-1.91 (m, 1H), 1.95-2.08 (dtt, J = 14.0, 12.8, 3.0 Hz, 1H), 2.09-2.17 (m, 1H), 2.60-2.73 (m, 2H), 3.03-3.13 (dt, J = 13.2, 3.0 Hz, 1H), 3.17-3.24 (td, J = 13.4, 3.7 Hz, 1H), 3.61-3.71 (ddd, J = 14.0, 12.0, 2.5 Hz, 1H) (**Fig. S4.**); ^13^C NMR (100 MHz, CDCl_3_, δ): 15.29, 23.47, 25.74, 51.92 (**Fig. S5.**). [M+H]^+^ was observed for C_4_H_8_OS_2_ and found 137.00914 Da (**Fig. S6.**) via Bruker 9.4T SolariX XR Mass Spectrometer (Bruker, Billerica, MA). Melting point found to be 83-86 °C (lit 83-86 °C).

### Expression and purification of *wild-type* SOD1, SOD1^A4V^, SOD1^G93A^, SOD1^H46R^, and SOD1^G85R^

Expression and purification of SOD1 were conducted as previously published (21). Briefly, EGy118ΔSOD1 yeast were transformed with a *wild-type* SOD1, SOD1^A4V^, SOD1^G93A^, SOD1^H46R^ or SOD1^G85R^ YEp351 expression vector and grown at 30 °C for 44-hr. Cultures were centrifuged, lysed in a blender using 0.5 mm glass beads, and subjected to a 60% ammonium sulfate precipitation. Then the sample was centrifuged, and the resulting supernatant was diluted to 2.0 M ammonium sulfate. The diluted sample was passed through a phenyl-sepharose 6 fast flow (high sub) hydrophobic interaction chromatography column (Cytiva Life Sciences, Marlborough, MA, USA) using a linearly decreasing salt gradient from high salt buffer (2.0 M ammonium sulfate, 50 mM potassium phosphate dibasic, 150 mM sodium chloride, 0.1 mM EDTA, 0.25 mM DTT, pH 7.0) to low salt buffer (50 mM potassium phosphate dibasic, 150 mM sodium chloride, 0.1 mM EDTA, 0.25 mM DTT, pH 7.0) over 300 mL. Fractions containing SOD1 eluted between 1.6 and 1.1 M ammonium sulfate and were confirmed with SDS-PAGE. These fractions were pooled and exchanged into low salt buffer (10 mM Tris pH 8.0). Pooled fractions were then passed through a Mono Q 10/100 anion exchange column (Cytiva Life Sciences, Marlborough, MA, USA) using a linearly increasing salt gradient from low salt buffer to high salt buffer (10 mM Tris pH 8.0, 1 M sodium chloride) from 0 – 30%. SOD1 fractions were collected between 5 and 12% high salt buffer and were confirmed with SDS-PAGE, western blot, and Fourier Transform Ion Cyclotron Resonance Mass Spectrometry (FT-ICR-MS).

### Confirmation of cross-link formation in vitro

*Wild-type* SOD1, SOD1^A4V^, SOD1^G93A^, SOD1^H46R^, and SOD1^G85R^ stock solutions were diluted to 40 μM in 10 mM ammonium acetate, pH 7.4. DMSO stock of *S*-XL6 was used to prepare 400 μM (10x) in 10 mM ammonium acetate (0.5% DMSO). Protein and cross-linker samples were combined in equal volumes (final concentration 20 μM SOD1, 200 μM cross-linkers, 0.25% DMSO) and incubated at 37 °C for 4 hours at 350 rpm. Complete cross-linking was confirmed by mass spectrometry on a 9.4T Bruker SolariX (Bruker Corporation, Billerica, MA) via direct infusion as previously described (25, 43). Prior to infusion, samples were diluted to 1 μM in 50:50 acetonitrile:water, with 0.1% formic acid. During analysis, 32 scans were acquired in positive mode and averaged. Funnel 1 and skimmer 1 were kept around 150 V and 20 V, respectively, and funnel RF amplitude was held at 60.0 Vpp.

### Differential Scanning Fluorimetry

The effect of cross-link formation on the thermal stability of SOD1 was determined by differential scanning fluorimetry as previously described (21, 27). *Wild-type* SOD1, SOD1^A4V^, SOD1^G93A^, SOD1^H46R^, and SOD1^G85R^ (20 μM in 10 mM ammonium acetate, pH 7.4) were incubated with ten-fold molar excess *S*-XL6 for 4 hours at 37 °C in protein low bind Eppendorf tubes using Eppendorf Thermomixer at 350 rpm (Eppendorf North America, Enfield, CT, USA). After incubation, SYPRO Orange (Invitrogen Corporation, Carlsbad, California, USA), an environmentally sensitive fluorescent dye which is quenched in an aqueous environment but becomes unquenched once it binds hydrophobic residues, was added to the reaction mixture to a final concentration of 200X. The samples were then transferred to a fast optical 96-well reaction plate (Applied Biosystem, Life Technologies Corporation, Carlsbad, California, USA) and loaded on to the real-time PCR machine (Applied Biosystem, Life Technologies Corporation, Carlsbad, California, USA) after plate was spun down to eliminate bubbles. The Melt curve template in StepOne Plus software was used to set up a method. The SYBR reporter and ROX quencher data were collected for 35 cycles with a temperature gradient ranging from 25 to 99.9°C at a ramp rate of 0.5°C/min. The final concentration of DMSO in the reaction mixture was checked as it attributes to the denaturation of proteins. Dilutions were performed accordingly to maintain the final concentration of DMSO less than 1% (vol/vol). All samples were run in triplicate. The background data was subtracted from the average relative fluorescence and average derivate for each temperature point, and the data were normalized before T_u_s were quantified by averaging the negative first derivative of relative fluorescence of all three runs and identifying the local minima.

### Hep G2 cell culture

The Hep G2 human hepatocarcinoma cell line (American Type Culture Collection, ATCC, Manassas, VA) was cultured using EMEM (Eagle’s minimal essential medium, ATCC) supplemented with 10% FBS (fetal bovine serum, ATCC) and 1% penicillin streptomycin as previously described (31) (10,000 units/mL penicillin and 10,000 μg/mL streptomycin, Fisher scientific, Hampton, NH). The cells were grown and sub-cultured around 72 hours at 37°C, 5% CO_2_ under controlled humidity. The cells growth and morphology were inspected using an inverted Carl Zeiss microscope (Carl Zeiss Microscopy LLC, White Plains, NY).

### Cell viability assay/cell cytotoxicity assay

Using manufacturer’s instructions, Hep G2 cells were cultured for 24 hours with a density of 2×10^5^ cells/mL in a 96-well culture plate at 37 °C, 5% CO_2_ under controlled humidity. The cells were treated with *S*-XL6 ranging with 10, 100, 150, 250, 400, 600, 800, and 1000 μM for 24 hours (0.1% DMSO). Cell cytotoxicity was performed via the MTT (3-(4,5-dimethylthiazol-2-yl)-2,5-diphenyltetrazolium bromide) assay at 570 nm absorbance using BioTek synergy H1 (Vermont, USA) plate reader according to previously published paper (31). The MTT stock was made at 5 mg/mL using 0.9% sodium chloride solution which was purchased from Sigma Aldrich (St. Louis, MO). Twenty-four hours after dosing the plate, the cultured medium was aspirated and replaced with the equal volume of MTT solution (10X dilution using EMEM complete media). The plate was incubated for 4 hours (37 °C, 5% CO_2_). The MTT solution was replaced by the equal amount of acidified isopropanol (0.3% hydrochloric acid (v/v)) and then formed formazan was dissolved by shaking the plate gently for 30 minutes. The absorbance was measured at 570 nm for both the samples and acidified isopropanol for background calculation. Chlorpromazine and cells with 0.1% DMSO were used as positive control and vehicle, respectively. The cell viability of each concentrations was measured by normalizing against mean value of cells with 0.1% DMSO (vehicle).

### SDS-Page and western blot

The Hep G2 human hepatocarcinoma cell line (American Type Culture Collection, ATCC, Manassas, VA) was cultured using both PBS buffer only and complete EMEM (Eagle’s minimal essential medium, ATCC) media including 10% FBS (fetal bovine serum, ATCC) and 1% penicillin streptomycin (10,000 units/mL penicillin and 10,000 μg/mL streptomycin, Fisher scientific, Hampton, NH). Cells were counted and checked for morphology using an inverted Carl Zeiss microscope (Carl Zeiss Microscopy LLC, White Plains, NY). A 96-well plate (Corning Life Sciences, Tewksbury, MA) was used for culturing cells at 0.2 mill/mL concentration for 24-hours at 37°C, 5% CO_2_ under controlled humidity for western blot experiment. Cells (both PBS buffer only and completed EMEM media) were dosed with *S*-XL6 in triplicate at different concentrations (1 μM, 10 μM, and 100 μM) and incubated for 30-minutes. In house purified *wild-type* SOD1 was used as positive control (standard) in the experiment. Existing buffer was aspirated and replaced with 30 μL of lysis buffer (150mM NaCl, 1% Triton X-100, 0.5% sodium deoxycholate, 0.1% SDS, and 50mM Tris, pH 8.0, all chemicals were purchased from Sigma Aldrich (St. Louis, MO)) and incubated for 10-minutes followed by addition of non-reducing sample buffer. Samples were loaded into a Bio-Rad mini TGX gel (Bio-Rad Life Sciences, Hercules, CA) along with PageRuler™ prestained protein ladder (ThermoFisher Scientific, Waltham, MA) and ran using a Bio-Rad electrophoresis cell. Carefully the gel was removed and transferred into a beaker filled with transfer buffer (25mM Tris, 192mM glycine, 0.1% SDS, and 10mM β-mercaptoethanol, all chemicals were purchased from Sigma Aldrich (St. Louis, MO)) and incubated briefly at 90°C and then transferred to a Trans-Blot Turbo mini nitrocellulose membrane using Trans-Blot Turbo Transfer system (Bio-Rad, Hercules, CA). The membrane was incubated overnight in antibody buffer (50mM tris-HCl, 150mM sodium chloride, 0.1% tween-20, 2.5% dry milk, all chemicals were purchased from Sigma Aldrich (St. Louis, MO)) with Cu/Zn SOD polyclonal antibody (1:1000 dilution, Enzo Life Sciences, Farmingdale, NY). Next day, the membrane was incubated with HRP-linked antibody buffer (1:1000 dilution, Cell Signaling Technology, Danvers, MA). The membrane was incubated briefly in dark with enhanced chemiluminescent (ECL, Thermo Fisher Scientific, Waltham, MA), followed by imaging using BioRad Chemidoc MP Imaging System (Bio-Rad, Hercules, CA).

### Mouse dosing and SOD1 isolation

Pharmacodynamic profiling was carried out using hemizygous mice expressing human SOD1^G93A^ (“fast-line” Jackson Laboratory; B6SJL-Tg(SOD1*G93A)1Gur/J, also known as SOD1-G93A stock-002726) (13). Mice were bred to express YFP in neurons in anticipation of BBB-penetration assays and Matrix-Assisted Laser Desorption Ionization Mass Spectrometry Imaging (MALDI MSI) analysis. Extraction of SOD1^G93A^ from red blood cell (RBC) protein was performed as previously described (32). *S*-XL6 was prepared in 1x PBS and mice were dosed at 10 mg/kg via intravenous (IV) injection in their lateral tail vein slowly and gently. Mice were trapped in restrainer and around 40 μL of blood was collected in Greiner Bio-one K_3_EDTA tubes (Greiner Bio-one, North Carolina, USA) post injection at 30-min, 1 hour, 2 hour, 4 hour, 8 hour, 12 hour, 24 hour, 48 hour, 72 hour, and 168 hour. Tubes were centrifuged at 2000 rpm at 4 °C for 5 minutes immediately after collecting blood. Carefully plasma was discarded and acid citrate dextrose solution (0.48% citric acid, 1.32% sodium citrate, 1.47% glucose, all chemicals were purchased from Sigma Aldrich (St. Louis, MO)) was used to wash the sample by centrifuging at 2000 rpm at 4 °C for 5 minutes. Following wash, the supernatant was removed, and RBC were lysed by the addition of 8x equivalent of 10 mM ammonium acetate. To the hemolysate, 0.15 equivalents of cold chloroform and 0.25 equivalents of cold ethanol was added. Samples were vortexed at 1800 rpm at 4 °C for 15 minutes and centrifuged at 12000 rpm for 10 minutes. The supernatant was collected and stored at −80 °C after flash freezing for LC-MS analysis. Prior to LC-MS analysis, samples were acidified to 10% formic acid. This study was performed in accordance with the Guide for the Care and Use of Laboratory Animals (National Institutes of Health, Bethesda, MD, USA). The protocol for this experiment with mice was approved by the Northeastern University Institutional Animal Care and Use Committee (IACUC).

### Confirmation of *S*-XL6 cross-linked SOD1^G93A^ dimer formation in vivo

In vivo crosslinking from purified RBC was confirmed using an H-Class Acquity UPLC (Ultra Performance Liquid Chromatography) system coupled to a Xevo G2-S Q-ToF (Quadrupole Time of Flight) mass spectrometer (Waters Corp, Milford, MA) as previously described (44, 45). The LC system was equipped with reversed phase Acquity UPLC Protein BEH C4 (300 Å pore size, 1.7 μm particle size, 100 mm bed length, 2.1 mm ID × 100 mm) column at 60 °C with a flow rate of 0.2 mL/min. The mobile phase consisted of a mixture of 0.1% formic acid in water (solvent A) and 0.1% formic acid in acetonitrile (solvent B). The sample was introduced in 10% formic acid and 5 μL was injected for analysis. UNIFI software (Waters Corp, Milford, MA) was used for system control and data processing. Solvents A and B were combined in a gradient: 0-2 min: 95% A; 2-70 min: 30% A; 72-75 min: 5% A; 78-80 min: return to initial conditions. The MS was operated in positive mode and calibrated prior to analysis (*ca.* daily). The chromatographic window containing the SOD1^G93A^ monomer and dimer were assigned using EICs (Extracted Ion Chromatogram) *m/z* 1322.5-1323.5 (monomer) and *m/z* 1180.0-1180.5 (dimer) and mass spectra from this region were summed. Raw MS data from this composite spectrum were deconvoluted and average masses were calculated using the MaxEnt1 algorithm. The ratio of dimer intensity (31836 Da, SD 1.3 Da) to the sum of monomer (15858 Da, SD 0.8 Da) plus dimer intensity were then used to calculate the percentage of SOD1^G93A^ dimer at different time points. 10 picomole of the SOD1^G93A^ monomer and dimer were analyzed (as above) in individual experiments and their MS signal intensities were within 10% (i.e., the differences in their chromatographic retention and ionization efficiency were within experimental error). The half-life of the *S*-XL6 cross-linked complex was calculated from apparent half-life of disappearance of the % dimer by linear regression of the terminal beta phase (24 to 168 hours time point).

### Small-angle x-ray solution scattering (SAXS)

SAXS data were collected at the G1 beamline at Cornell High Energy Synchrotron Source (CHESS). For each sample, two measurements were taken, one of the protein with buffer and one with buffer by itself. Solution scattering data were captured every second for 10 frames. The 10 frames of both buffer and protein were then averaged and the buffer was subtracted out to get the scattering for the protein. Samples were run in a 96-well plate and held at 4 °C continuously. Data collection was in the scattering angle (*q*) range of 0.008 to 0.71 Å^−1^ and processed using the software, RAW.

### Preparation of SOD1 samples for SAXS analysis

SOD1^G93A^ was prepared at 3 mg/mL (approx. 190 μM) in HEPES buffer (115 mM NaCl, 1.2 mM CaCl_2_, 1.2 mM MgCl_2_, 2.4 mM K_2_HPO_4_, 20 mM HEPES, pH 7.4). Stock solutions of *S*-XL6 was freshly made at 10 mM in HPLC grade methanol and diluted to 946 μM in HPLC grade water (five-fold concentration of protein). 100 μL of 3 mg/mL protein was mixed with 110 μL of 946 μM of *S*-XL6. Control sample contained 5% MeOH (final conc. 2.5%). Samples were then incubated at 37 °C for 6 hours to ensure complete cross-linking. After incubation, excess compound was buffer exchanged out of each sample using a 10 kDa MWCO ultrafiltration device. 320 μL is diluted to 15 mL into HEPES buffer, spun down to approximately 500 μL, and resuspended in another 15 mL of HEPES buffer. After the final spin, samples were removed and concentrated in a smaller ultrafiltration device and brought to approximately 70 μL final volume (4.7 μM). Samples were flash frozen and stored at –80 °C prior to analysis.

### SAXS data analysis and reconstruction of molecular envelopes

Programs within the ATSAS suite (46) were used to determine the estimated radius of gyration and three-dimensional molecular envelopes for SOD1^G93A^ with and without *S*-XL6 using the x-ray solution scattering data. The GNOM program was used to evaluate the pair distribution plot using an indirect Fourier transform. SOD1^G93A^ with *S*-XL6 had a *D*_max_ value of ~79 Å whereas untreated SOD1^G93A^ had a much higher *D*_max_ of over 200 Å due to protein unfolding. The GASBOR program was used to generate three-dimensional *ab initio* models of connected beads to fit the GNOM data, with the number of beads set approximately to the total number of amino acids in the SOD1 constructs. In order to assess the uniqueness of these solutions, 10 bead models were generated without any symmetry applied, then compared and averaged. Figures were produced using CHIMERA (47) followed by superimposition of envelopes.

### Peptide-level hydrogen deuterium exchange mass spectrometry (H/D-X MS)

To 5ul of 40μM SOD1 sample (in 10mM Ammonium Acetate in H_2_O, pH 7.4) was added 20μl of 99% D_2_O sample buffer (10mM ammonium acetate in D_2_O, pH 7.3), diluting the concentration of the protein down to 8μM. For the reference sample, HPLC grade water was added instead of D_2_O buffer. Samples with D_2_O buffer were incubated at 37 °C for five exposure timepoints: 15 seconds, 50 seconds, 500 seconds, 1hr, and 4hrs. All exchange reactions occurred at 37 °C, pH ~7.4 to mimic in vivo conditions. The reaction mixture was quenched by the addition of 25μL of quench buffer (8M Gunadinium Hydrochloride (GnHCL), 0.5M tris(2-carboxyethyl) phosphine (TCEP), 0.2M Citric Acid at pH 2.35), lowering the pH of the final mixture down to pH 2.45, diluting the concentration of protein in the sample down to 4μM while simultaneously denaturing the protein and reducing the disulfides. For the 15 second timepoint, to ensure timely quenching of the exchange reaction, D_2_O buffer previously stored at 37 °C was added and the incubation itself was performed at room temperature. All samples preparations were performed in triplicates, flash frozen immediately after the quenching reaction and stored at −70 °C until analysis. Prior to analysis, 50μl of 0.1% Formic Acid in H_2_O was added to the sample to reduce the GnHCL concentration down to 2M, the recommended concentration threshold for the pepsin column. This was immediately followed by injecting the sample onto a Waters UPLC system designed for H/D-X MS analysis where the samples were digested, desalted, and separated, online. The digestion and trapping of peptides was carried out during a 3 minute trapping step over an immobilized pepsin column with a flow rate of 100μl/min in 0.1% formic acid and water at 10 °C. The peptides were trapped on an ACQUITY HSS T3 100 Å, 1.8 μM trap column (Waters Corp, Milford, MA) maintained at 0 °C. At the end of the trapping step, within the 0 °C chamber, the flow was directed to the ACQUITY HSS T3 100Å, 1.8 μM analytical column (Waters Corp, Milford, MA) at 75μl/min (average back pressure was around 7500 psi). The analytical separation step was performed over a 9min gradient of 5-25% (0-7min) of buffer B; 25-95% (7-8.5min) of buffer B (buffer A, 0.1% formic acid in water, buffer B, 0.1% formic acid in acetonitrile). Eluate from the analytical column was directed into a Waters QToF Synapt G2 HD mass spectrometer with electrospray ionization and lock-mass correction (using the Glu-fibrinogen peptide). Blanks (data not shown) were used between each sample injection to ensure there is no carryover of peptides between runs. Mass spectra were acquired between 50 – 2000 m/z, in positive polarity and resolution mode. Scan time was set to 0.5 seconds, cone voltage to 30V, capillary was 3.5kV, trap collision energy was 6V and desolvation temperature of 175 °C.

Prior to H/D-X MS experiments, sample preparation and run conditions were optimized by varying concentrations of GnHCL (2M, 4M, 8M), incubation temperature with D_2_O buffer (4 °C, 37 °C), flow rate over the pepsin column and HPLC gradient. Conditions listed above yielded the best sequence coverage (greater than 98%) and resolution in the shortest run time. Furthermore, to minimize rate of back exchange from D back to H, the sample mixture pH after the quench reaction was lowered to approximately 2.5 and majority of the run on the instrument was performed at 0 °C (48). All H/D-X MS experiments were performed under identical conditions, therefore deuterium levels reported are relative and were not corrected for back exchange. Optimized sample preparation and instrument conditions to minimize back exchange from deuterium to hydrogen and triplicate measurements for each sample allowed for high confidence comparability assessments between sample types (49).

### H/D-X MS data analysis

Mass spectra and chromatography data were acquired using MassLynx (Waters Corp). The peptides were identified and confirmed via MS^2^ using the PLGS software. The PLGS generated peptide lists and MassLynx acquired mass spectra were imported into the DynamX™ HDX data analysis software 3.0 (Waters Corp) for further analysis. Only peptides that were identified across all six replicates of each SOD1 variant (example: triplicates of untreated *wild-type* SOD1 and *S*-XL6 cross-linked *wild-type* SOD1) were considered. The maximum sequence length of peptides was set to 45, minimum peak intensity set to 10,000, and maximum MH+ error set to 10ppm. Sequence coverage across all samples achieved was ~99% and the average redundancy for covered amino acids was 11. The mass spectra were processed within the DynamX software by centroiding the isotopic distribution of various charge states for all the peptides (typically +2,+3, +4). Deuterium uptake levels were measured by calculating the differences between the centroid of the deuterated peptide vs. the undeuterated reference peptide. These mass shifts and differences are plotted against the exchange timepoints (**Fig. S3** for the SOD1^A4V^, SOD1^H46R^, SOD1^G85R^ N and C terminal peptides). Final percentage uptake was calculated by averaging the percent uptake across triplicate measurements utilizing overlapping peptides and recurring residues to elucidate residue level uptake measurements, wherever possible. Three replicates of each sample type were run on different days, and the standard deviation was ~3% or less. A conservative threshold of 5% was set during the comparative analysis between sample types and only consistent differences in % deuterium uptake above 5% have been emphasized. Peptic maps were obtained from the DynamX software. Maestro 11.8 (Schrödinger Maestro, New York, USA) was used to map the conformational changes onto the crystal structure of *wild-type* SOD1 (PDB: 1SPD).

### Determination of Cu and Zn metal content

All samples were sent to Element Materials Technology (Santa Fe Springs, CA) for quantitation of Cu and Zn metal content via Inductively Coupled Plasma Mass Spectrometry (50). Briefly, a sample portion (0.05 g) was mixed with internal standards (In-Tb-Sc) and then diluted to a final mass of 5 g with a solution of 0.1% ammonium hydroxide, 0.05% EDTA, and 0.05% Triton X100. The sample appeared to have completely dissolved. Elements were analyzed on an Agilent 7500 ICP-MS (Agilent Technologies) with an octopole collision cell. The standard operating conditions used were RF power: 1550W, Sample depth: 8mm, Carrier gas flow: 1L/min, Spray chamber temperature: 2 °C, Nebulizer Pump: 0.1 rps, Collision gas: 5.4 mL/min helium. For quality control, spike recovery experiments were performed where detection limits of Cu and Zn were measured to be 0.01 ppm and 1 ppm, respectively. Metal content for untreated and S-XL6 treated samples are reported in **Table S2**.

### Proteolytic digestion and MALDI-TOF-MS peptide analysis

Samples of SOD1^A4V^ with or without ten-fold molar excess of *S*-XL6 in 10 mM Tris HCl, pH 7.4 were incubated for 4 hours at 37 °C (51). After incubation, samples were alkylated with iodoacetamide (100 mM for 30 minutes), heated to 75 °C for 20 minutes, and then treated with two volumes of Poroszyme immobilized trypsin (Applied Biosystems, Life Technologies Corporation, Carlsbad, CA, USA) at 37 °C for 15 minutes, mixing every few minutes to keep beads suspended. Beads were removed by centrifugation before analysis. SOD1^H46R^ was incubated with or without ten-fold molar excess of *S*-XL6 and deuterated *S*-XL6 followed by pepsin digestion (1:20 w/w, pepsin:protein) for 120 minutes. Both the digested samples were analyzed using a microflex MALDI-TOF mass spectrometer (Bruker Daltonics, Billerica, Massachusetts, USA) in reflectron mode in the 2-5 kDa range and linear mode in the 4-20 kDa range. Spectra were calibrated using Peptide and Protein I Calibrant (Bruker Daltonics, Billerica, Massachusetts, USA). Matrix only and trypsin digest reaction mixture without SOD1 spectra were acquired as negative controls. Spectra were analyzed in flexAnalysis and BioTools 3.2 (Bruker Daltonics, Billerica, Massachusetts, USA). Peptide mass fingerprinting was performed using MASCOT (Matrix Science, Boston, MA, USA) using trypsin as the enzyme with up to 5 missed cleavages, 100 ppm mass tolerance, and cysteine carbamidomethylation as a variable modification.

## Supporting information

Supplemental Section

## Acknowledgements

We thank Dr. George Bou-Assaf for their guidance with the H/D-X MS experiments, Luis Viskatis and Dr. Brian Fahie for their support and encouragement.

