## Supplemental Section for "Protein crosslinking as a therapeutic strategy for SOD1-related ALS"

**This PDF file includes:**

Figures S1 to S6
Tables S1 and S2

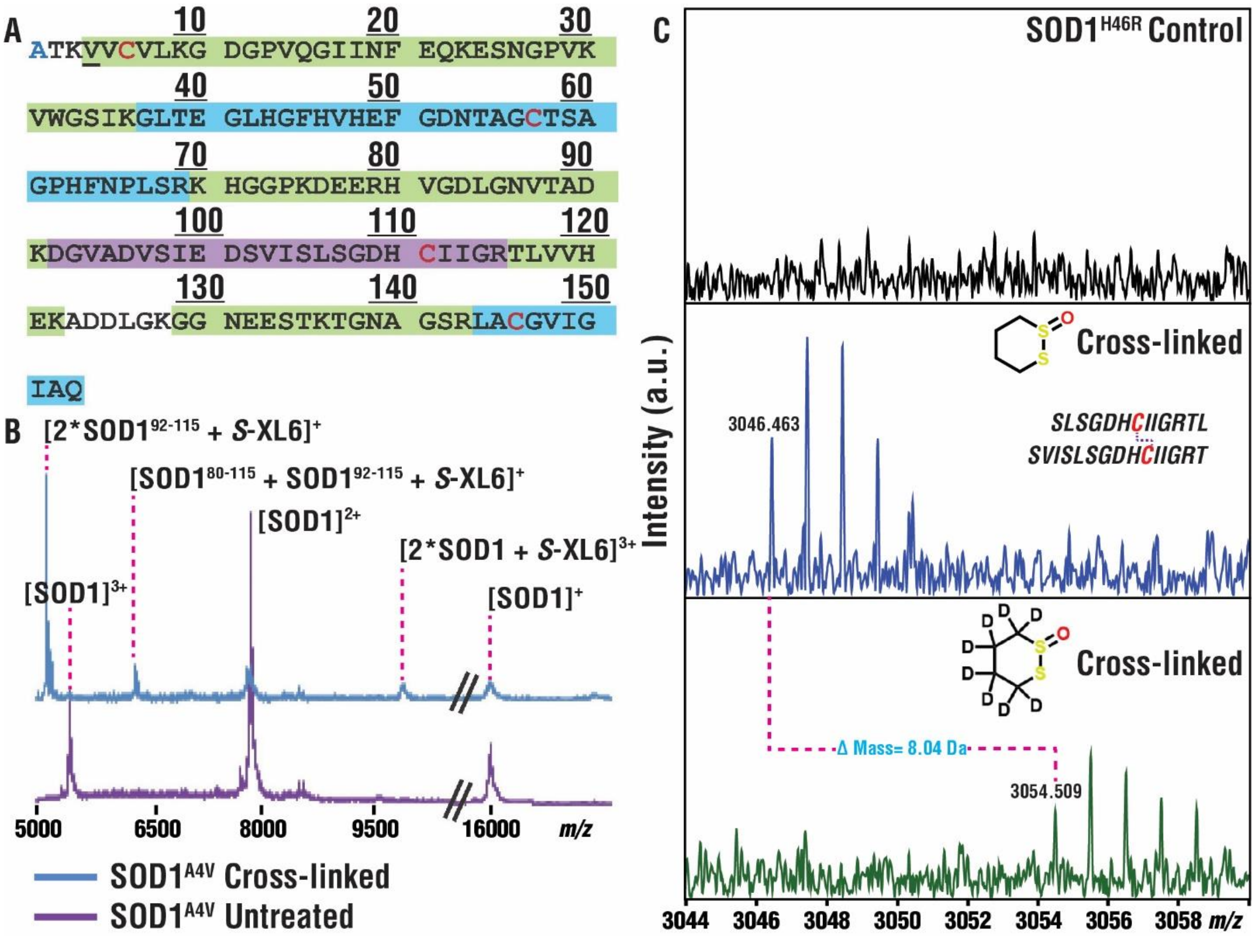


**Fig. S1**. *S*-XL6 cross-links SOD1 at Cys111. (*A*) Trypsin digest and MALDI-ToF-MS analysis of SOD1$^{A4V}$ gave a combined sequence coverage of 94%, including peaks corresponding to the Cys57-Cys146 disulfide linked peptides (blue) and the Cys111- containing peptide (purple) linked by *S*-XL6. N-terminal acetylation is shown in blue text, cysteines are shown in red text, SOD1$^{A4V}$ mutation is shown underlined. (*B*) Examining the higher mass range revealed multiply charged forms of SOD1$^{A4V}$ including the *S*-XL6 linked dimer, as well as two different peaks corresponding to Cys111-linked peptides. (*C*) MALDI-FTICR-MS analysis of pepsin digested SOD1$^{H46R}$. Top panel shows SOD1$^{H46R}$ control sample and mid panel exhibits peaks corresponding to Cys111 linked peptides via *S*-XL6. The linked peptides are shown in black text, and cysteines are highlighted in red that are crosslinked. The bottom panel represents same crosslinked peptides with deuterated *S*-XL6 demonstrating a mass shift of 8.04 Da confirming the selectivity of the crosslinker to Cys111 residue (mechanism of action of CTs).

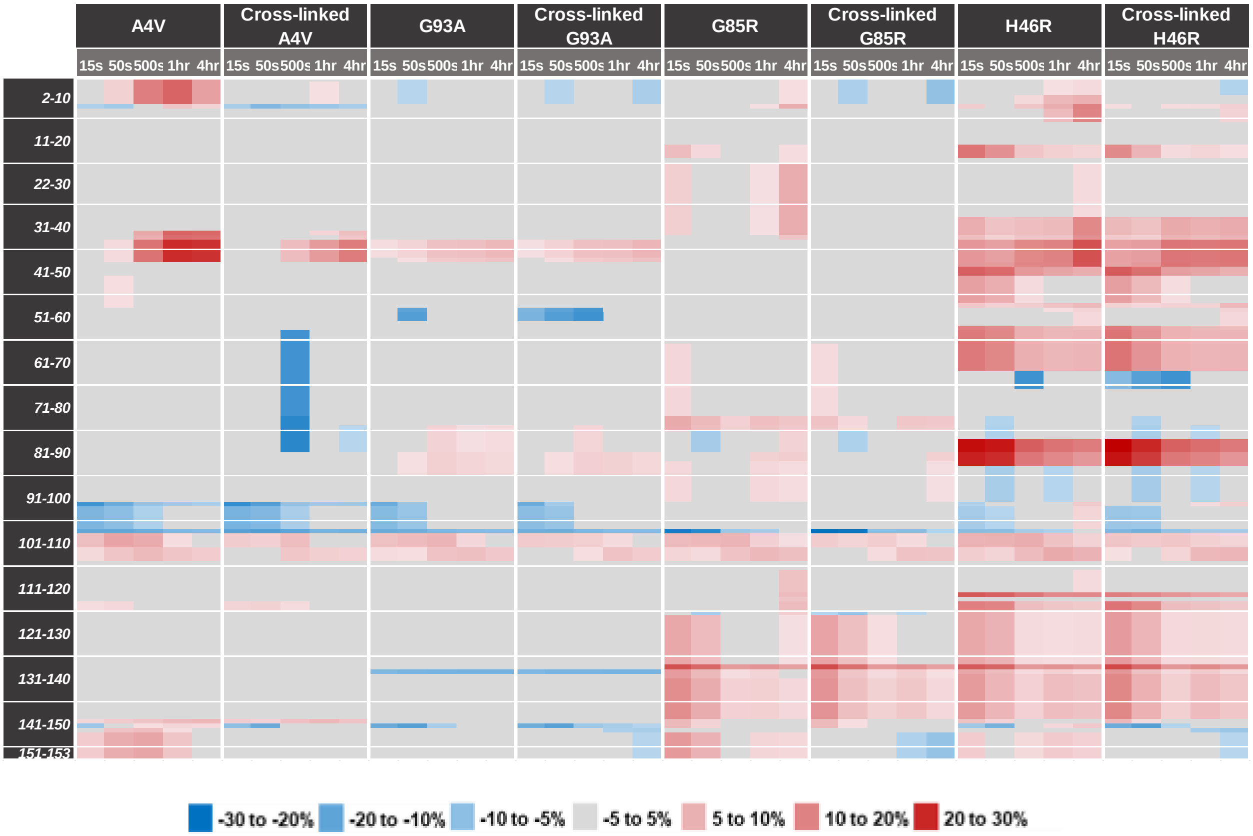


**Fig. S2. *S*-XL6 cross-linking of SOD1 variants regulates the structure making them more like the stable, dimeric *wild-type* form**. Differences in deuterium uptake (ΔU, legend shown below) of untreated and crosslinked variants for all timepoints (15s, 50s, 500s, 1 hour, 4 hours) compared to the stable dimeric *wild-type* SOD1 are reported here. For the 4 hr timepoint the prominent ΔU values are as follows: ***N-terminus and nearby:*** *residues 2-7* SOD1$^{A4V}$ untreated 13.8% à 1.2% cross-linked; *residues 2-7* SOD1$^{G93A}$ untreated 3.4% à -6.2% cross-linked; *residues 2-7* SOD1$^{G85R}$ untreated 5.3% à -8.4% cross-linked, *residue 8* SOD1$^{G85R}$ untreated 11.9% à -1.8% cross-linked; *residues 6-7* SOD1$^{H46R}$ untreated 11.5% à 0.6% cross-linked, *residues 9-11* SOD1$^{H46R}$ untreated 17.7% à 6.8% cross-linked. ***Residues 39-43 and nearby****:* SOD1$^{A4V}$ untreated 29% à 17.6% cross-linked. *Residues 22-37,* SOD1$^{G85R}$ untreated 11.9% à 2.9% cross-liked; SOD1$^{H46R}$ untreated 24.6% à 19.7% cross-linked*.* ***Residues 111-116****:* SOD1$^{G85R}$ untreated 9.2% à -2.0% cross-linked*.* ***C-terminus****:* SOD1$^{G85R}$ untreated 6.2% à-8.1% cross-linked; SOD1$^{H46R}$ untreated 7.1% à -5.1% cross-linked.

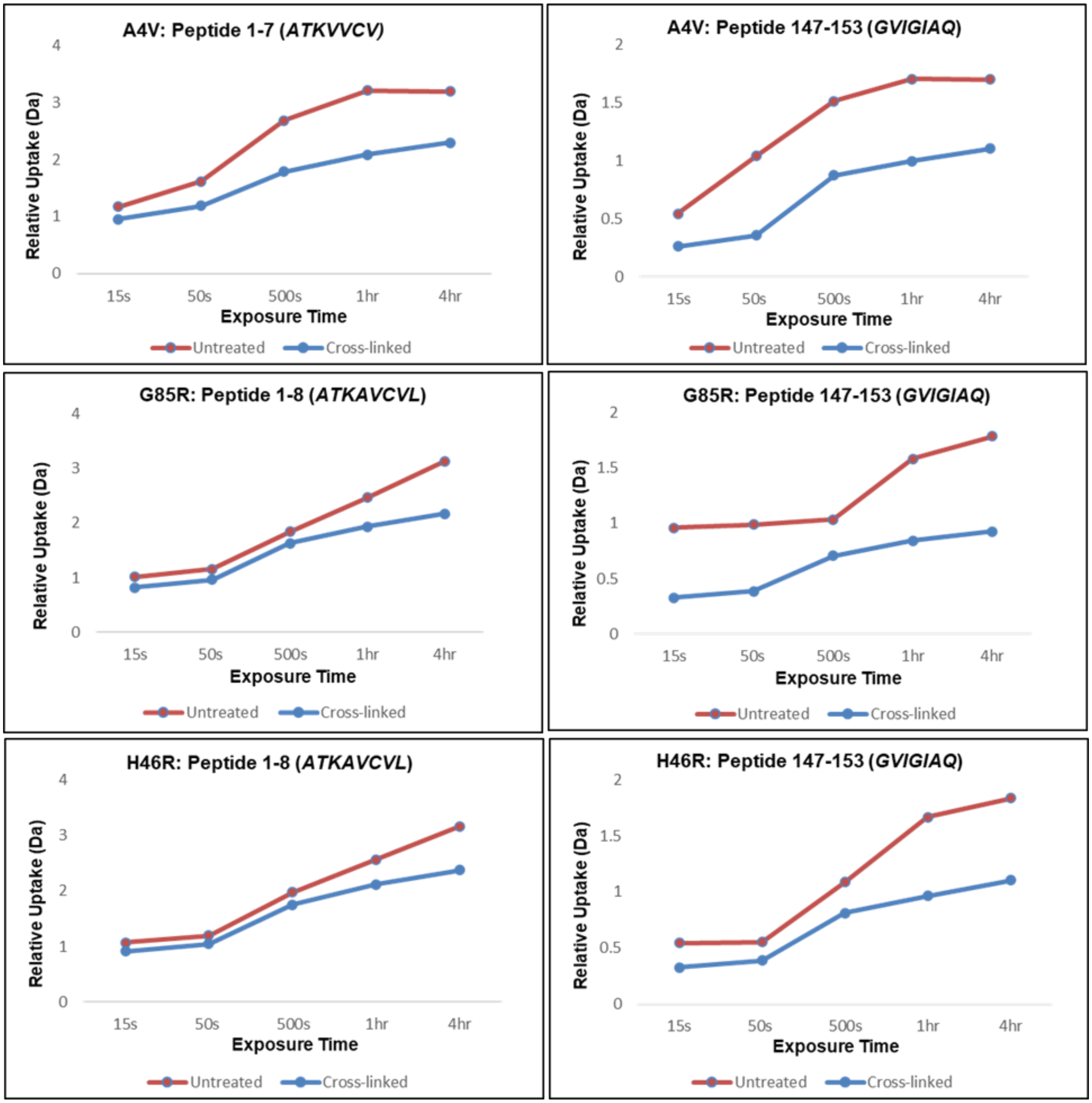


**Fig. S3**: ***S*-XL6 has stabilizing structural effects on SOD1 fALS variants**. Uptake plots for the non-comparable terminal peptides for $SOD1^{A4V}$, $SOD1^{G85R}$, $SOD1^{H46R}$ are shown. Exposure timepoints (15s, 50s, 500s, 1hr, 4hr) are represented on the x-axis plotted against the relative deuterium uptake (Da) on the y-axis. Red lines represent the untreated samples and blue lines represent the *S*-XL6 cross-linked samples. Uptake is generally higher for the control samples, and differences continue to grow with later timepoints (1hr, 4hr).

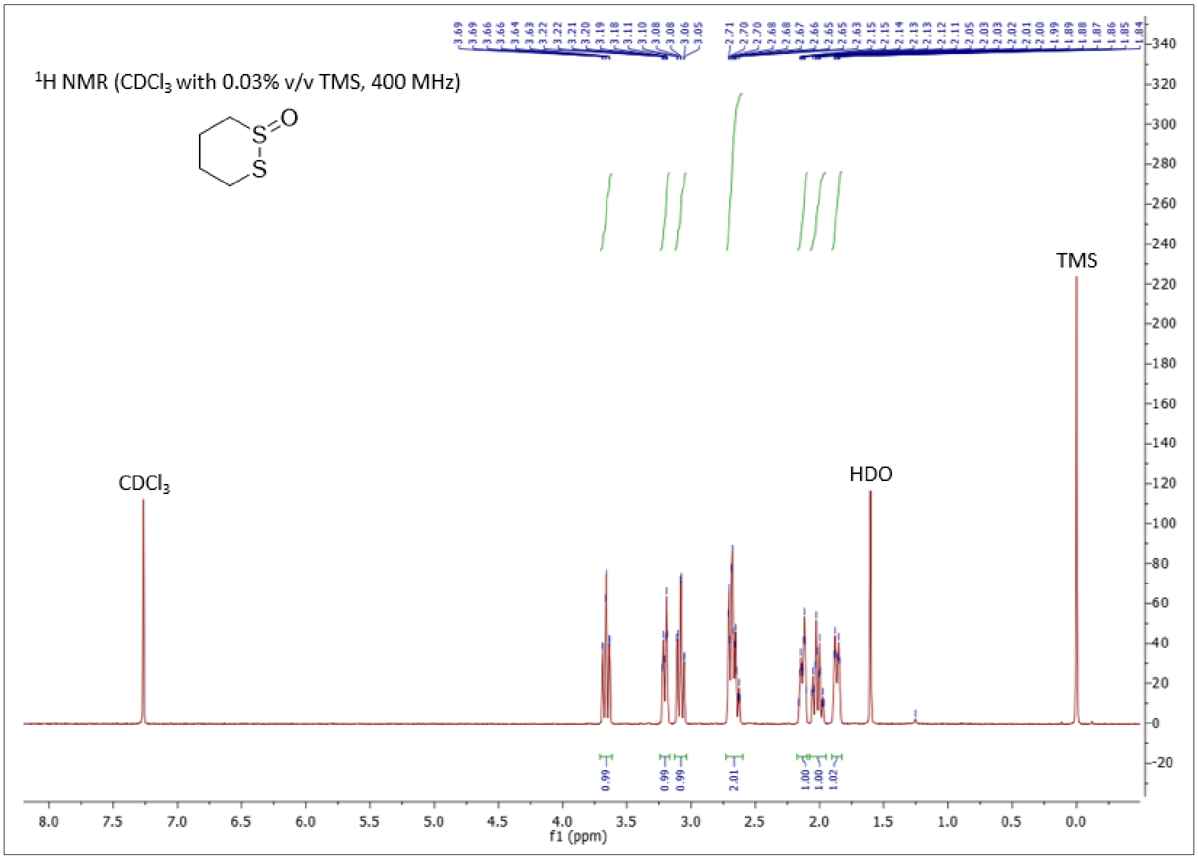


**Fig. S4.** $^1$H NMR of *S*-XL6.

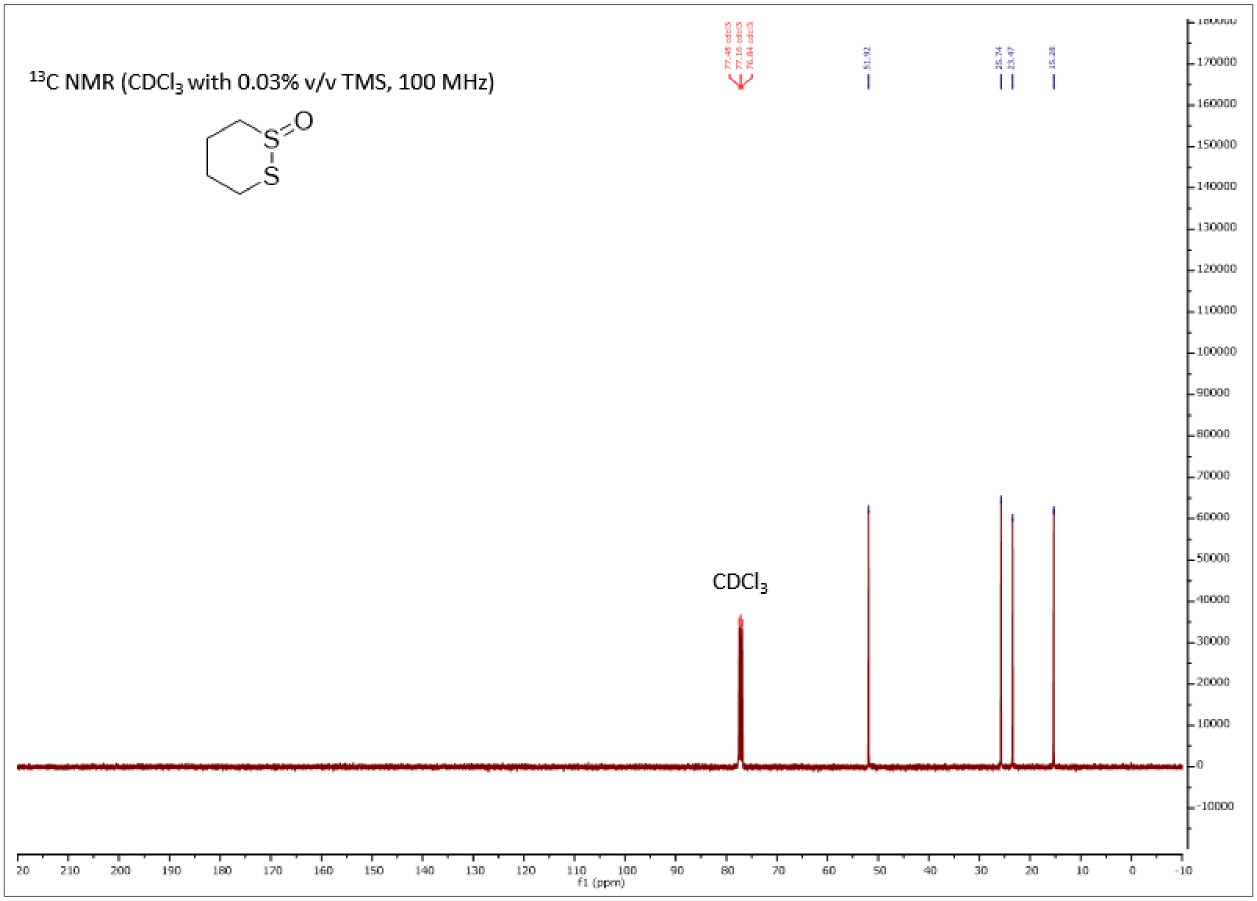


**Fig. S5.** $^{13}$C NMR of *S*-XL6.

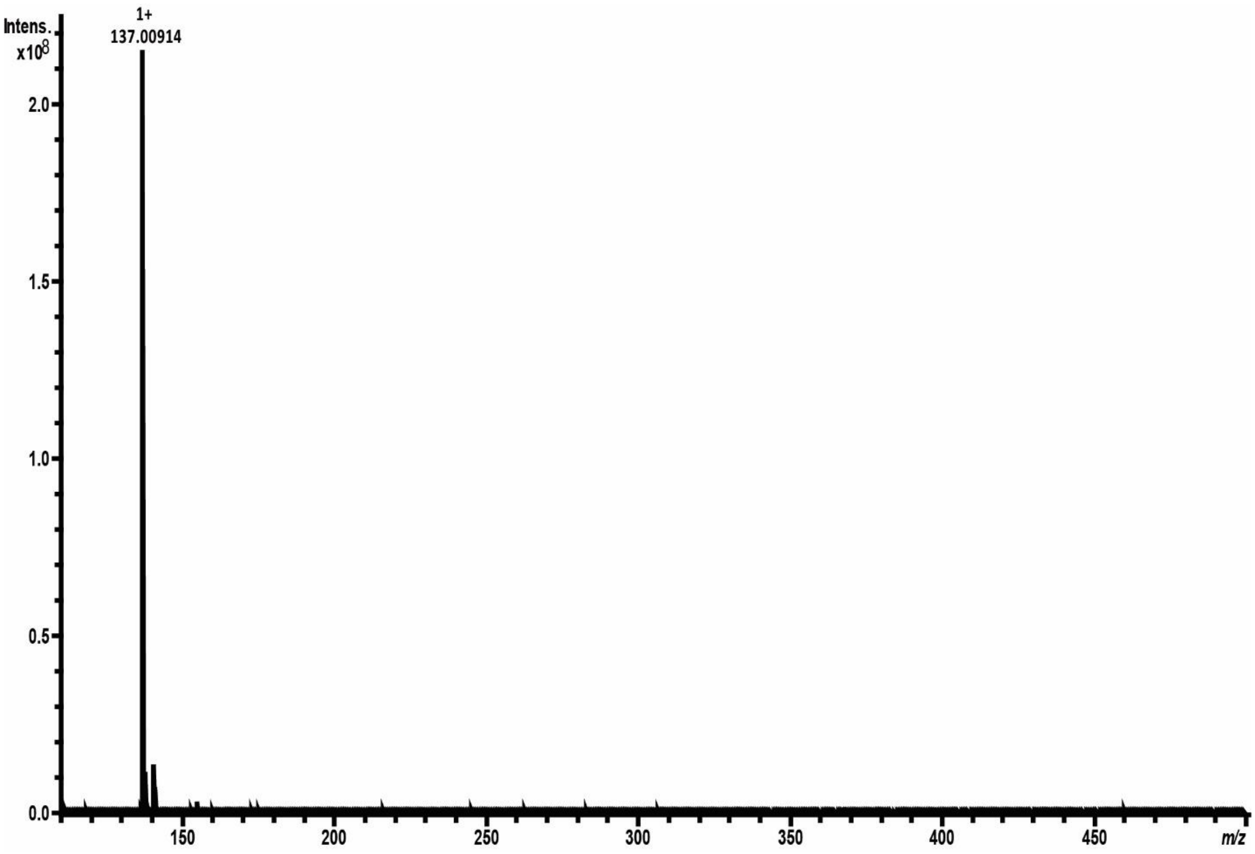


**Fig. S6.** High resolution mass spectrum of *S*-XL6 showing observed $[M+H]^+$ mass 137.00914 Da.

| Time (hr.) | % Dimer |
|---|---|
| **0.5** | **38** |
| **1** | **63** |
| **2** | **55** |
| **4** | **53** |
| **8** | **51** |
| **12** | **42** |
| **24** | **26** |
| **48** | **20** |
| **72** | **16** |
| **168** | **6** |

**Table S1.** Pharmacodynamic analysis of *S*-XL6 cross-linked $SOD1^{G93A}$ dimer. Blood from $SOD1^{G93A}$ mice was collected at different time points post *S*-XL6 dosing at 10 mg/kg via tail vein injection. Table shows the percentage of $SOD1^{G93A}$ cross-linked dimer at the different timepoints.

<table>
<tr><th>SOD1</th><th>Cu (ppm)</th><th>Zn (ppm)</th><th>Cu:SOD1</th><th>Zn:SOD1</th></tr>
<tr><td>WT</td><td>1.10</td><td>6</td><td>0.46</td><td>2.43</td></tr>
<tr><td>Cross-linked WT</td><td>1.20</td><td>5</td><td>0.50</td><td>2.03</td></tr>
<tr><td>A4V</td><td>0.68</td><td>6</td><td>0.28</td><td>2.43</td></tr>
<tr><td>Cross-linked A4V</td><td>0.74</td><td>6</td><td>0.31</td><td>2.43</td></tr>
<tr><td>G93A</td><td>1.20</td><td>5</td><td>0.50</td><td>2.03</td></tr>
<tr><td>Cross-linked G93A</td><td>1.30</td><td>5</td><td>0.54</td><td>2.03</td></tr>
<tr><td>G85R</td><td>0.04</td><td>ND</td><td>0.02</td><td rowspan="4">N/A</td></tr>
<tr><td>Cross-linked G85R</td><td>0.05</td><td>ND</td><td>0.02</td></tr>
<tr><td>H46R</td><td>ND</td><td>ND</td><td rowspan="2">N/A</td></tr>
<tr><td>Cross-linked H46R</td><td>ND</td><td>ND</td></tr>
</table>

**Table S2. *S*-XL6 does not alter the metal binding affinity for SOD1 variants.** Utilizing ICP-MS, we confirmed the metal content for each SOD1 sample: *wild-type*-like variants ($SOD1^{A4V}$ and $SOD1^{G93A}$) were partially metallated and metal deficient variants ($SOD1^{G85R}$ and $SOD1^{H46R}$) contained little to no metal**.** Metal content is calculated per monomer. The detection limit for Copper is 0.02 ppm and Zinc 2 ppm. ND: not detected.
